# Regulation of major bacterial survival strategies by transcript sequestration in a membraneless organelle

**DOI:** 10.1101/2022.01.06.475198

**Authors:** Tamar Szoke, Nitsan Albocher, Omer Goldberger, Meshi Barsheshet, Anat Nussbaum-Shochat, Reuven Wiener, Maya Schuldiner, Orna Amster-Choder

## Abstract

Liquid-liquid phase separation (LLPS) of proteins was shown in recent years to regulate spatial organization of cell content without the need for membrane encapsulation in eukaryotes and prokaryotes. Yet evidence for the relevance of LLPS for bacterial cell functionality is largely missing. Here we show that the sugar metabolism-regulating clusters, recently shown by us to assemble in the *E. coli* cell poles by means of the novel protein TmaR, are formed via LLPS. A mutant screen uncovered residues and motifs in TmaR that are important for its condensation. Upon overexpression, TmaR undergoes irreversible liquid-to-solid transition, similar to the transition of disease-causing proteins in human, which impairs bacterial cell morphology and proliferation. Not only does RNA contribute to TmaR phase separation, but by ensuring polar localization and stability of flagella-related transcripts, TmaR enables cell motility and biofilm formation, thus providing a linkage between LLPS and major survival strategies in bacteria.

## INTRODUCTION

*E. coli* cells lack membrane-bounded organelles. Still these cells respond rapidly to environmental changes via pathways that could benefit from compartmentalization of specialized domains. The ability of both protein and RNA molecules to localize to specific domains within the bacterial cell has been intensively studied in recent years (*1*–*5*). Cases showing that membraneless organelles may form in bacteria via liquid–liquid phase separation (LLPS) have been reported in the last couple of years (*6*). Examples thus far include DNA segregation (*7*), cell division proteins (*8*), *E. coli* RNA polymerase (*9*), *Caulobacter crescentus* RNA degradosome, a signaling microdomain components at the poles of the asymmetrically dividing *C. crescentus* (*10, 11*) and an ABC transporter in Mycobacterium tuberculosis (*12*). Yet, the direct relevance of the physical state of these phase-separated proteins for their function remains obscure.

Data accumulated in recent years revealed that, despite the lack of membrane-bounded organelles, bacterial cells have a higher level of spatial organization than originally assumed. Specifically, the poles of *E. coli* cells are emerging as hubs for sensing and responding, but the mechanisms involved in assembling macromolecules there are largely unknown. We recently showed that TmaR (previously called YeeX) clusters to one pole in *E. coli* cells when phosphorylated on a tyrosine. We could also demonstrate that this ability is important for cell survival in mild acidic conditions, which *E. coli* often encounters in its natural habitats (*13*). We have further shown that TmaR controls the activity of the general protein of the phosphotransferase system (PTS), Enzyme I (EI), the main regulator of sugar utilization in most bacteria, by its polar sequestration and release (*13*). The dynamics of TmaR, mainly within the pole region, the lack of evidence for its membrane-anchoring, and the rapid change between clustered and diffused EI (*14*), intrigued us to posit that TmaR forms polar condensates, which compartmentalize EI, via LLPS.

Here, we explored the mechanism that underlies formation of polar clusters by TmaR and its effect on cell physiology and functionality. By characterizing its liquid-like behavior, we show that TmaR forms condensates at the cell poles, and that purified TmaR forms droplets *in vitro*. To uncover motifs that affect TmaR condensation, we performed a high content screen of a library of strains each expressing a TmaR mutant with a single base substitution and isolated TmaR variants that fail to undergo LLPS. We show that upon overexpression, TmaR forms filamentous structures, reminiscent of the irreversible liquid-to-solid transition of disease-causing proteins in eukaryotes (*15*), which may extend from cell to cell and affect cell division. Our results reveal an unexpected reciprocity between TmaR and RNA molecules. On one hand, RNA contributes to the formation of TmaR condensates. On the other hand, only when in condensates, TmaR protects *fliA* and *motA* transcripts, thus controlling the motility, swarming and biofilm formation capabilities of *E. coli*, providing a new role for TmaR and for LLPS in bacteria.

## RESULTS

### TmaR clusters that control sugar utilization assemble trough liquid-liquid phase-separation

To explore the mechanism underlying TmaR cluster formation, we monitored the behavior of endogenously expressed mYFP-TmaR clusters over time by time-lapse microscopy. We detected small clusters of TmaR merging into bigger clusters, whereas other clusters were observed splitting into smaller ones (**Fig. 1A**), a characteristic behavior reported for proteins that undergo LLPS in eukaryotes (*16*) and recently also in bacteria (*6*). Since liquid condensates formed by LLPS were shown to recover within minutes from photobleaching (*17*), we used fluorescence recovery after photobleaching (FRAP) to further validate that TmaR clusters exhibit the dynamics expected from proteins undergoing phase separation. The graph in **Fig. 1B** shows that a considerable amount of fluorescent TmaR molecules entered the polar microdomain in less than a minute following photobleaching, and additional molecules continued to join the cluster for the rest of the experiment (see images at the bottom of **Fig. 1B** that show a cluster in a representative cell before bleaching and during the 5 minutes of recovery). Another tool commonly used for characterizing condensates formed by LLPS is the aliphatic alcohol 1,6-hexanediol, shown to disassemble phase-separating liquid condensates, but not solid structures (*18*). The images in **Fig. 1C** show complete dispersal of a TmaR cluster in a representative cell treated with 10% 1,6-hexanediol, as opposed to clusters in untreated cells (mock experiment, **Fig. S1A**, right). It is worth mentioning that condensate dispersal upon addition of the 1,6-hexanediol was very fast, occurring in a timescale of seconds, and could be observed already at the earliest time point (time 0), taken immediately after addition of the hexanediol (**Fig. S1A**, left). Together, the timescale of TmaR cluster dynamics and their 1,6-hexanediol-induced dispersal suggest that they are formed by LLPS.

**Fig. 1.**
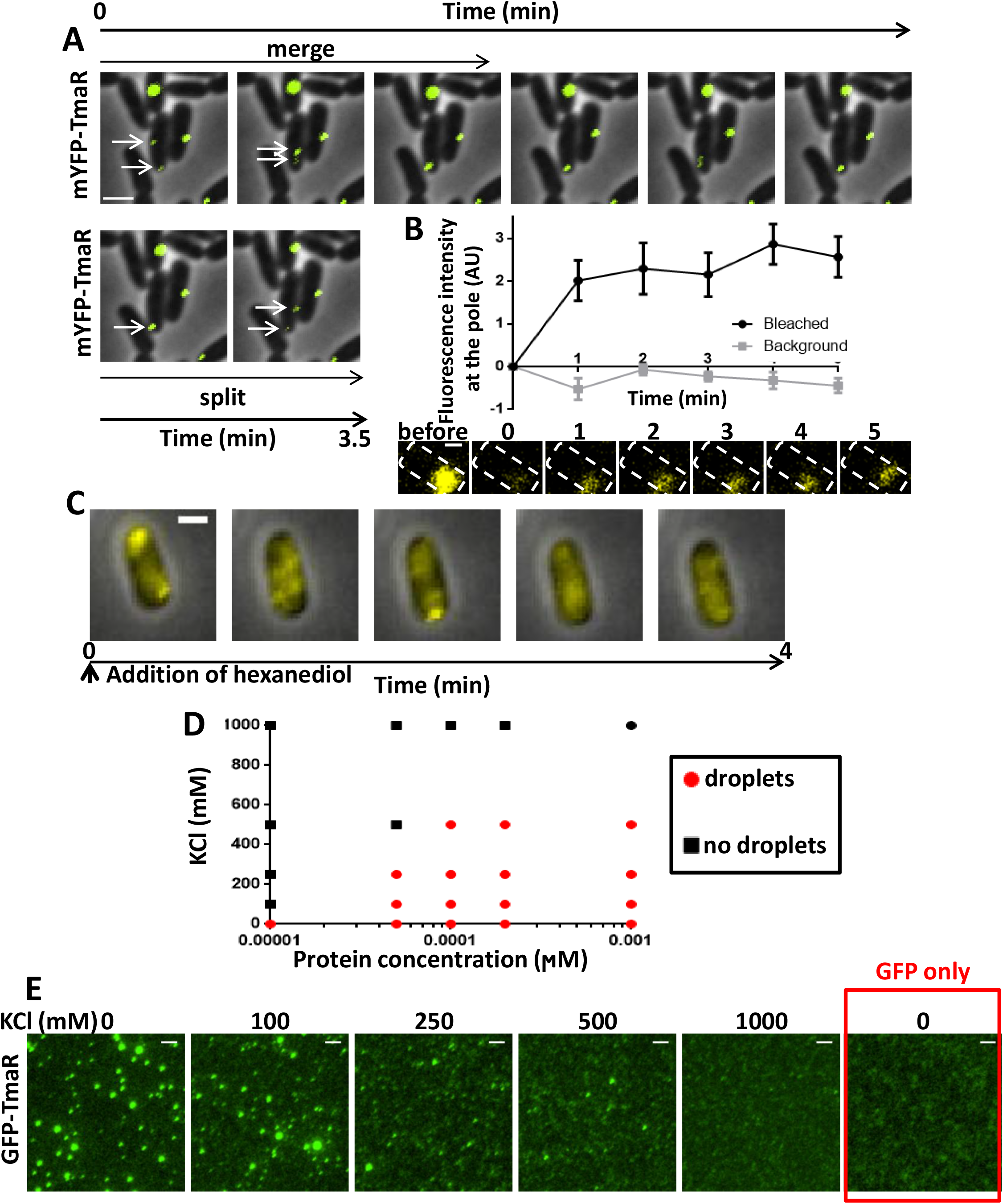
TmaR forms condensates in *E. coli* cells and droplets *in vitro* by LLPS. (A) Images of cells expressing TmaR tagged with monomeric YFP (mYFP-TmaR, yellow) from *tmaR* chromosomal locus obtained by time-lapse microscopy. Cells were spotted on an agar pad and pictures were taken every 30 second for 3.5 minutes. Scale bar, 2 µm. (B) Recovery following targeted photobleaching of TmaR-YFP condensates in *E. coli* cells by laser scanning confocal microscopy. **Upper panel:** Line chart showing the recovery dynamics of the YFP signal in the bleached region (Bleached, black) compared to the signal in a non-bleached region (Background, gray), immediately after bleaching (time 0) and every minute for 5 minutes. Mean and standard errors are shown (n=9). **Lower panel:** FRAP images of a representative cell before bleaching and every minute for 5 min after bleaching. A dashed line marks the outline of the cell. Scale bar, 0.5 µm. (C) Images of a representative cell endogenously expressing mYFP-TmaR obtained by time-lapse microscopy. Cells were spotted on an agar pad containing 10% 1,6-hexanediol and pictures were taken every minute for 4 min. Scale bar, 1 µm. See Fig. S1A for additional cells and for images of mock experiment. (D) Phase diagram of purified GFP-TmaR incubated in 50 mM Tris at the indicated concentrations of protein and KCl. Black squares indicate conditions in which the protein is dissolved (no droplets); red circles indicate conditions in which the protein is condensed (droplets). See fluorescence microscopy results in Fig. S3A. (E) Images of purified GFP-TmaR, at a final concentration of 0.001 µM, incubated in 50 mM Tris with the indicated concentrations of KCl. The GFP only control, at a final concentration of 1 µM, is in 50 mM Tris buffer with no salt. Scale bar, 2 µm.

EI, the major sugar utilization regulator, was shown to localize with the TmaR clusters and to be activated by its release from them, but not to cluster on its own or affect TmaR clustering (*13*). Hence, we investigated its dynamics in time-lapse microscopy and its response to hexanediol treatment. The results in **Fig. S1B** show that EI-mCherry clusters in the same cells presented in **Fig. 1A** colocalize with one of the TmaR clusters that merges with other clusters and splits. Similar to TmaR, EI clusters dissolve immediately upon adding 10% hexanediol to the cells (**Fig. S1C**, left panels), as opposed clusters in untreated cells (mock experiment, **Fig. S1C**, right panels). Hence, as expected from a membraneless constituent, which is not capable of phase separating on its own, EI takes after TmaR.

We next aimed at assessing whether TmaR phase separation depends on additional cellular factors or whether it can phase-separate on its own, as reported for many proteins (*17*). To address this issue, we purified GFP-tagged TmaR and asked whether it can form phase separated droplets and under which conditions. A common feature of proteins that undergo LLPS is that they form droplets outside the cell in a concentration- and salt-dependent manner (*17*). To see whether this is true for TmaR, we visualized purified GFP-TmaR at different concentrations in Tris buffer containing different KCl concentrations by fluorescence microscopy. As shown in Fig. **S1D**, the purified protein formed droplets under the conditions depicted by red dots in the phase diagram presented in **Fig. 1D**. In the absence of KCl, GFP-TmaR formed droplets in all the protein concentrations tested, whereas in 1M KCl, no droplets were observed and the GFP signal was spread uniformly. Droplets were also observed with purified mCherry-tagged TmaR (**Fig. S1E**) Hence, TmaR can phase separate on its own *in vitro* and the process depends on protein and salt concentration.

Looking more carefully at the salt-dependence of purified GFP-TmaR revealed that it shifts from a dispersed to a condensed state between 100 mM and 250 mM KCl (**Fig. 1E**), indicating that phase separation of the purified protein occurs under physiological conditions (*17*). Of note, purified GFP at the same concentration does not form droplets even in the absence of KCl (**Fig. 1E**, right image), demonstrating that the TmaR moiety in GFP-TmaR, and not the GFP tag, trigger LLPS. Measuring the droplets area revealed that upon increasing the salt concentration, the clusters size is reduced till a minimal size is reached at 500 mM KCl (**Fig. S1F**, left graph). The number of droplets, on the other hand, stayed constant until 250 mM KCl and sharply dropped at higher salt concentrations (**Fig. S1F**, right graph). Together, these results indicate that TmaR forms droplets upon reaching a certain threshold level, but the size of the condensates, and supposedly the amount of protein in them is affected by the concentration of salt

Another factor that might affect droplet formation is the time of incubation (*17*). Indeed, the total area of droplets, measured in several fields, at the start of purified TmaR incubation and after 3 or 5 h of incubation increased constantly with time (**Fig. S1G**).

Crowding agents are known to promote droplet formation by phase separating proteins, as they increase their factual concentration (*8*). In line with that, addition of PEG 8000 to a final concentration of 10% resulted in an increase in TmaR-GFP droplets size (**Fig. S1H**), reinforcing the dependence of TmaR phase separation on its concentration. To test the degree of fluidity of purified GFP-TmaR in the droplets, we used FRAP. The observed fast reformation of the droplets after photobleaching (**Fig. S1I)** is in the time scale expected for phase separating proteins. Together, the behavior of purified TmaR *in vitro* supports the notion that TmaR is capable of undergoing LLPS by itself.

### Identification of residues and motifs that are required for TmaR condensates formation

The compliance of TmaR with the many criteria for a protein that undergoes phase separation provided us with an opportunity to learn more about the requirements for phase separation in bacteria and in general. We first asked whether TmaR contains known motifs and regions often associated with LLPS. Since intrinsically disordered protein regions (IDRs) are known as drivers of LLPS (*17*), we checked if TmaR is predicted to contain such regions, using PrDOS (http://prdos.hgc.jp/cgi-bin/top.cgi). The prediction suggests that TmaR has a very short IDR at its N’-terminus, and a longer one at its C’-terminus **(Fig. S2A)**. We therefore constructed two chromosomal fusions between truncated *tmaR* and mYFP, one without the C’-terminal IDR (“no IDR”) and the other of only the C’-terminal IDR (“IDR only”). Notably, both fusion proteins did not form condensates (**Fig. 2A**), suggesting that the C’-terminal IDR of TmaR is necessary but not sufficient for LLPS. Except for the IDR, most of TmaR protein is predicted to fold into two alpha helices that form a coiled-coil structure (**Fig. S2B**). When each of the helices was fused to mYFP, neither construct formed condensates (**Fig. 2B**). Hence, each alpha helix is necessary but not sufficient for TmaR LLPS, suggesting that the coiled coil structure that assembles from both helices is important for phase separation.

**Fig. 2.**
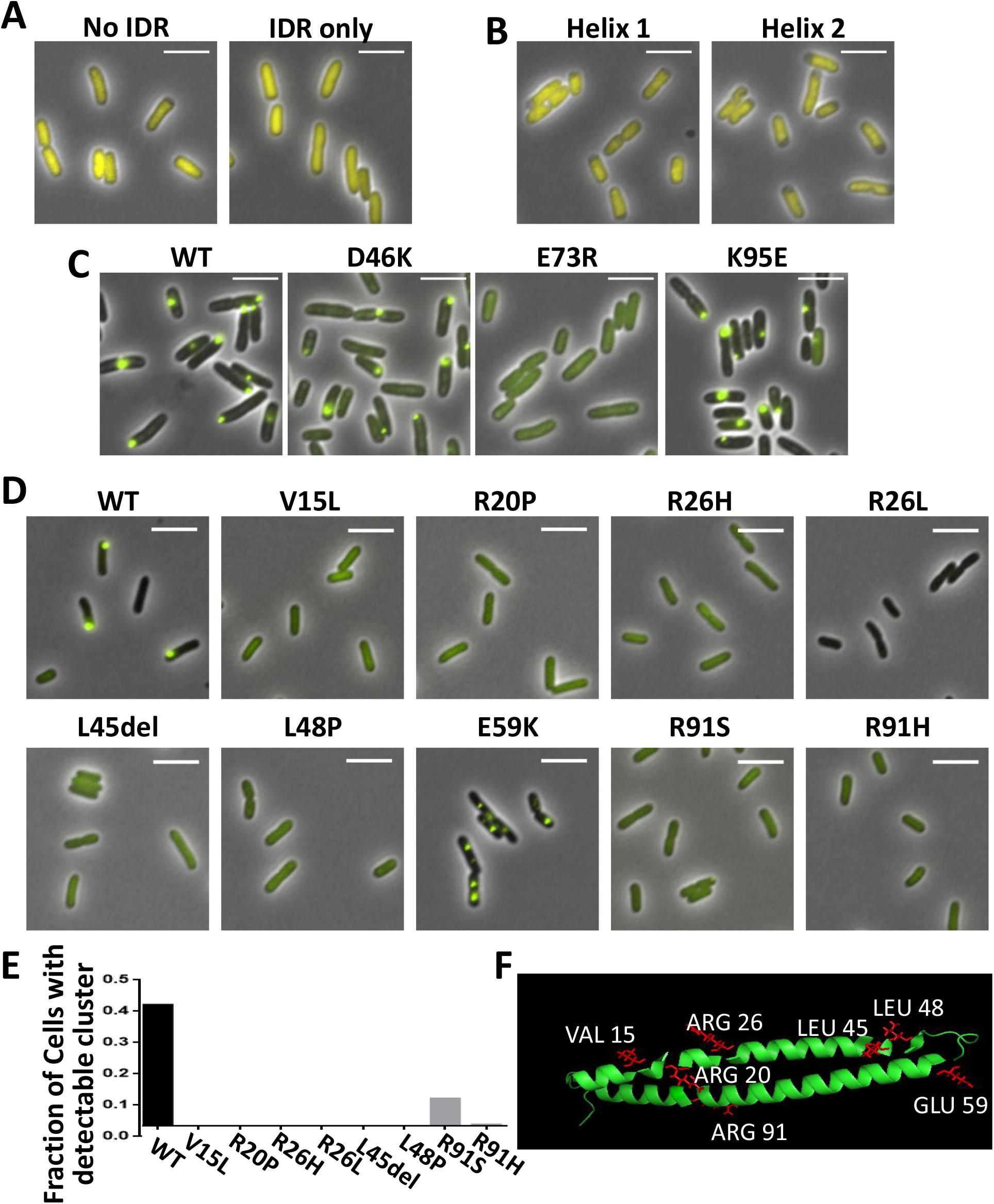
Identification of residues and motifs that are required for TmaR condensates formation by biased and unbiased approaches. (A) Images of cells in mid-logarithmic phase expressing mYFP-tagged truncated TmaR proteins lacking the IDR region (No IDR) or encompassing only the IDR region (IDR only), from the endogenous tmaR locus. Scale bar, 5 µm. (B) Images of cells expressing mYFP-tagged helix 1 or helix 2 of TmaR from *tmaR* endogenous locus and promoter. Scale bar, 5 µm. (C) Images of cells expressing YFP-tagged wild type TmaR (WT) or the indicated TmaR mutants from the *tmaR* endogenous locus. Scale bar, 5 µm. (D) Images of cells in mid-logarithmic phase expressing YFP-tagged wild type TmaR (WT) or the indicated TmaR mutants from the *tmaR* endogenous locus. L45 del denotes a missense mutation that resulted in the deletion of leucine in position 45. Scale bar, 5 µm. (E) The average fraction of cells with detectable clusters of TmaR-YFP in wild type (WT) or the indicated TmaR mutants. (F) TmaR structure prediction by trRosetta presented in PyMOL with the mutated residues found to change TmaR-YFP localization shown in red.

Weak electrostatic interactions, involving charged residues, often play a role in phase-separated condensate formation (*17*). Using EMBROSS.charge (https://www.bioinformatics.nl/cgi-bin/emboss/charge), we searched for regions in TmaR that are rich in charged residues and replaced three amino acids in these regions by amino acids with the opposite charge (**Fig S2C**, red arrows), thus constructing mYFP-TmaR D46K, mYFP-TmaR E73R and mYFP-TmaR K95E. While D46K and K95E did not affect TmaR clustering, the E73R mutation completely abolished TmaR clustering (**Fig. 2C**). Notably, we have previously shown that the neighboring residue of E73, a tyrosine in position 72 (Y72), gets phosphorylated, a modification often associated with LLPS (*19*), and that this event is required for TmaR activity as EI regulator (*13*). Jointly, the current and previous results highlight the importance of the negative patch around positions 96-77 in TmaR, and suggest that both the phosphorylation and the charge it confers affect TmaR ability to undergo LLPS *in vivo*.

Next, we aimed at identifying residues that affect TmaR capability to undergo LLPS by conducting an unbiased screen for TmaR mutants that fail to do so. To this end, we constructed a library of TmaR mutants by random mutagenesis (error prone PCR) of the *tmaR* gene fused to YFP and expressed from a low copy plasmid with its native promoter. The library was then introduced into cells deleted for the *tmaR* gene, and the subcellular distribution of the resulting mutant TmaR proteins was monitored by automated fluorescence microscopy. Out of the 768 mutants that were screened, we isolated eight TmaR mutants that failed to form clusters and were instead diffuse throughout the cytoplasm. The mutations were substitutions of six amino acids, V15, R20, R26, L45, L48 and R91 (**Fig. 2D, 2E and 2F**). The mutation in L45 was a missense mutation, which could influence the formation of the coiled coil, since L45 is predicted to be in close proximity to Y72, shown to be important for TmaR condensation (see above) in the 3D structure (**Figure S2D**). Two of the mutants, R20P and L48P, were an introduction of the helix breaking proline residue into the predicted alpha helix. Together, the two mutations reinforce the importance of the coiled coil structure for TmaR condensation.

In line with the reported importance of arginine for condensate formation (*20*), three of the six mutated residues were arginines, with two of them, R26 and R91, replaced by two different amino acids, histidine and leucine or histidine and serine, respectively. Of note, the replacement of arginine to leucine in position 26 showed a decrease in the overall YFP intensity, suggesting a lower stability of this mutant (**Fig. 2D**). The replacement of either of these two arginines by a histidine, which is also positively charged, together with replacement of valine in position 15 by leucine, both hydrophobic uncharged amino acids, suggest that it is not the charge of single amino acids that determines whether the protein phase separates. Rather, it is likely their structural constellation that enables the weak and promiscuous interactions required for phase separation.

Finally, our screen identified one mutant, a substitution of glutamic acid in position 59 by lysine, which forms clusters that do not localize to the poles (**Fig. 2D**). Since E59 is predicted to face out (**Fig. 2F**), the non-polar phenotype of the mutant suggests that this residue is involves in interactions with molecules that target TmaR condensates to the poles.

Overall, our screen identified the coiled-coil, the IDR and charged patches in TmaR as mutually important for its condensation via LLPS, and E59 as important for polar localization of its condensates.

### Liquid-to-solid state transition of TmaR affects cell morphology and physiology

In eukaryotes, failure to control condensate formation may cause transition to a non-native, irreversible - on biological timescales - and less dynamic solid-like state. The resulting aberrant insoluble structures are formed due to cellular stress, impairments in protein quality control, ageing-related loss of homeostasis or mutations and repeat-expansion disorders. These structures are often toxic to the cell and may lead to severe diseases (*21*–*24*). Although condensates formed by LLPS are high-order assemblies, they are still in a liquid state, whereas their misfolded structures, such as amyloids, prions, fibrilar structures and various other polymers and aggregates, have biophysical properties of gel-like or solid-like aggregates. Because proteins can fall out of solution when their concentration increases due to mis-regulated gene expression (*21*), e.g., disrupted transcriptional regulation during cancer development (*25*), we asked whether an increase in TmaR cellular concentration by its overexpression can induce such a transition. Notably, overexpression of TmaR led to the formation of chained cells in which TmaR was observed as a long filament/aggregate (**Fig. 3A**). Not only did TmaR form longer and more ordered filaments upon longer periods of overexpression (**Fig. 3A**, compare images I-III to images IV-V), but it was often observed at the borders of cells, sometimes accumulating in globular (**Fig. 3A**-I) or tubular (**Fig. 3A-**II & III) bodies that connect the cells. Using the aggregation predicting server PASTA 2.0 (http://old.protein.bio.unipd.it/pasta2/index.html), TmaR is predicted to contain a short sequence that causes aggregation (**Fig. S3A**).

**Fig. 3.**
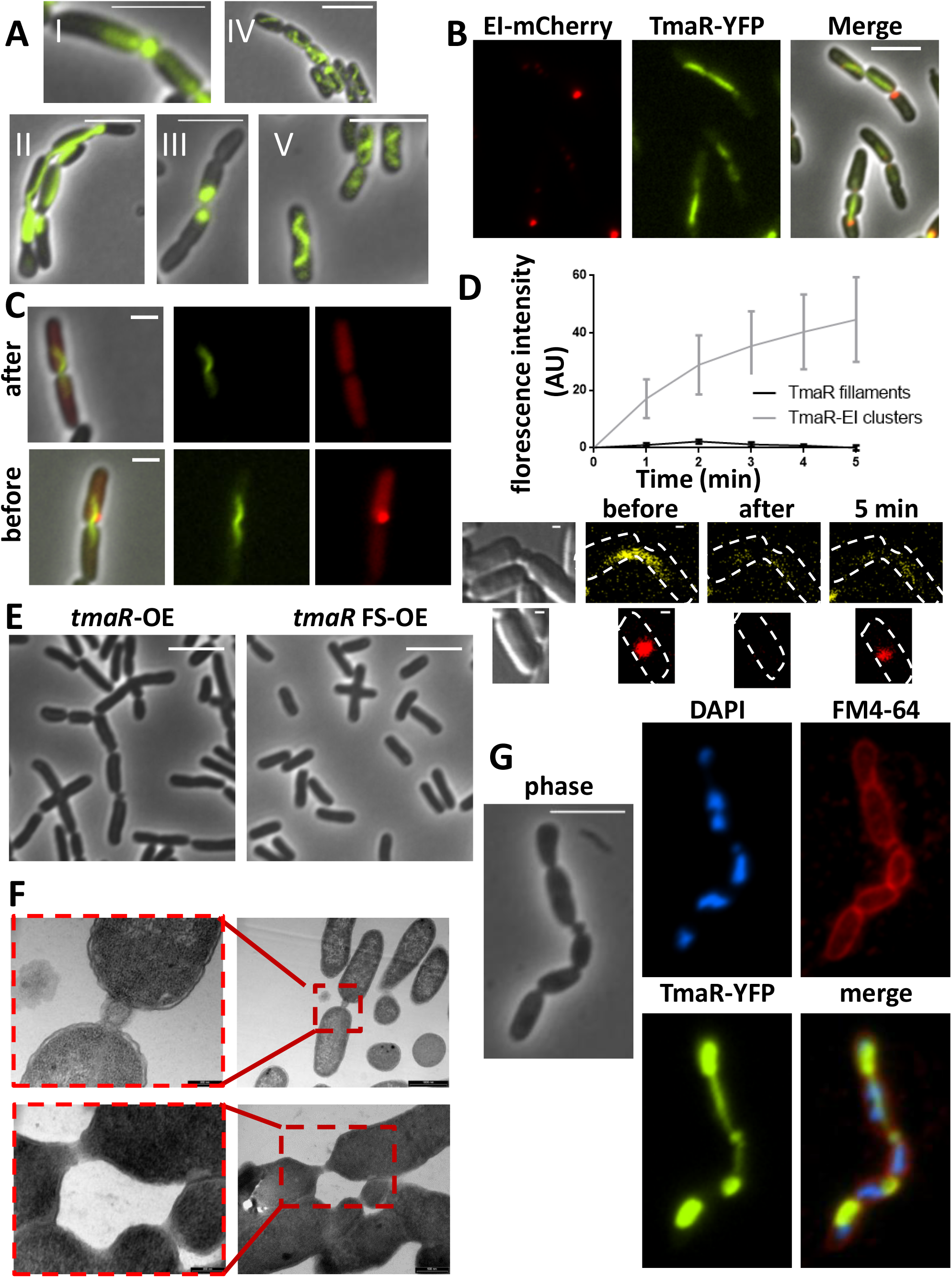
TmaR may undergo liquid-to-solid state transition that affects cell morphology and impairs cell division. (A) Images of representative cells overexpressing non-tagged TmaR from the pZE12 vector and endogenously expressing TmaR-YFP. Cells were grown for I-III: 1.5 h, IV-V: 2.5 h in the presence of 0.1 mM IPTG. Scale bar, 5 µM. (B) Images of cells overexpressing non-tagged TmaR from the pZE12 vector and endogenously expressing TmaR-YFP and EI-mCherry. Cells were grown in the presence of 0.1 mM IPTG. Scale bar, 5 µM. (C) Images of cells overexpressing non-tagged TmaR from the pZE12 vector and endogenously expressing TmaR-YFP and EI-mCherry obtained by time-lapse microscopy. Cells were grown with 0.1 mM IPTG and spotted on an agar pad containing 10% 1,6-hexanediol. Pictures were taken every 2 min for 10 min. Shown are the pictures taken at time 0 (before) and after 10 minutes (after). For images taken at the other time points see Fig. S5. Scale bar 2µM. (D) Recovery following targeted photobleaching of TmaR-YFP filaments and TmaR-YFP-EI-mCherry condensates in *E. coli* cells overexpressing non-tagged TmaR from the pZE12 vector and endogenously expressing TmaR-YFP and EI-mCherry. Cells were grown in the presence of 1 mM IPTG until mid-logarithmic phase and FRAP was performed by laser scanning confocal microscopy. **Upper panel:** Line chart showing the recovery dynamics of the YFP signal in the bleached TmaR filament (black) and in the TmaR-EI-mCherry condensate (gray), immediately after bleaching (time 0) and every minute for 5 minutes. Mean and standard errors are shown (n=9). **Lower panel:** FRAP images of a representative cells before bleaching (before), immediately after bleaching (after) and 5 min post bleaching (5 min). A dashed line marks the outline of the cell. Scale bar, 0.5 µm. (E) Images of cells overexpressing wild type *tmaR* (*tmaR*-OE) or *tmaR* with a frameshift mutation (*tmaR* FS-OE) from the pZE12 vector grown in the presence of 0.1 mM IPTG for 3 h. Scale bar, 5 µM. (F) Transmission electron micrographs of cells overexpressing TmaR. **Left**: TmaR expression was induced from the pZE12 vector with 0.1 mM IPTG for 3 h. Scale bar in lower magnitude picture, 1000 nm; scale bar in magnified image, 200 nm. **Right:** TmaR expression was induced from the pCA24N vector with 1 mM IPTG for 4h. Scale bar in lower magnitude images, 500 nm, and in the magnified images, 200 nm. Dashed boxes define the magnified areas. (G) Images of representative cells overexpressing non-tagged TmaR from the pZE12 vector and endogenously expressing TmaR-YFP. Cells were grown for 1.5 h and then the DNA was stained with DAPI and the membrane was stained with FM 4-64. Scale bar, 5 µM.

To learn about the consequences of TmaR transition upon an increase in its concentration, we monitored the distribution of EI, previously shown by us to be recruited to TmaR clusters, but not to form clusters on its own and not to affect TmaR clustering (*13*), in cells overexpressing TmaR. To this end, we co-expressed TmaR-YFP and EI-mCherry from their endogenous chromosomal loci in cells overexpressing non tagged TmaR. The results show that the majority of EI colocalized with the residual TmaR that remained in polar condensates and not with the bulk of the filamentous TmaR (**Fig. 3B**). This result suggested a scenario in which TmaR molecules in the polar condensates and in the filaments have different physical properties, that the two physical states of TmaR may co-exist in the cell, and that only the liquid state is recognized by EI. To test this assumption, we used 1,6-hexanediol to ask if TmaR aggregates are more solid than its polar condensates. Time-lapse microscopy of cells treated with 10% hexanediol demonstrated that the EI-mCherry-bound TmaR condensates dissolved, while the TmaR-YFP higher structures did not (**Fig. 3C** and **S3B**). Using FRAP, we verified that the two forms of TmaR have different solubility: while the recovery time after photobleaching of EI-mCherry-containing condensates was very fast, filamentous TmaR-YFP did not recover even after 5 minutes (**Fig. 3D**). Importantly, not only high concentration of TmaR can drive liquid-to-solid transition, but also a single amino acid substitution, since the results in **Fig. S3C** show that the clusters formed by the TmaR E59K mutant also do not dissolve in hexanediol. Of note, the cells expressing the TmaR E59K mutant also chain, but to a lesser extent than cells overexpressing WT TmaR (**Fig. S3D)**. Taken together, the results thus far show that TmaR is capable of transitioning from polar condensates to higher structures via a liquid-to-solid transition that is not reversible.

The phenomenon of cell chaining was not observed when overexpressing the TmaR-Y72F mutant, which does not phosphorylate and does not form phase-separated condensates (**Fig. S3E**). In line with our previous observation that upon its overexpression TmaR-Y72F forms inclusion bodies at the poles (*13*), overexpression of TmaR-Y72F formed polar aggregates that do not resemble the filamentous structures formed by wild type TmaR (**Fig. S3F**). Hence, not only is the phosphorylated tyrosine involved in LLPS, but it is also involved in the capacity to transition to a filamentous form.

The formation of chained cells upon TmaR overexpression suggested that the cells cannot complete division. Imaging GFP-tagged ZapA, a cell division protein that assists Z-ring formation in midcell, in cells overexpressing TmaR demonstrated that these cells are indeed defective in cell division, with ZapA distributing throughout the elongated cells, as opposed to the ZapA ring in midcell when TmaR in not overexpressed (**Fig. S3G**, compare TmaR-OE and empty vector images). Overexpression of *tmaR* with a frameshift mutation from the same plasmid did not produce such a phenomenon **(Fig. 3F**, right panel), implying that the high concentration of the TmaR protein is responsible for the defect in cell division. Cell division was impaired upon overexpression of TmaR from additional vectors that express TmaR at different levels, although to different degrees, often leading to the formation of extremely elongated cells (**Fig. S3H**). Observing the division site between the chained cells, formed upon TmaR overexpression, using transmission electron microscopy (TEM) validated that the chained cells are often connected by globular or tubular bodies (**Fig. 3F**), a phenomenon not observed with cells containing the vector only (**Fig. S3I**, right panel and insert), or cells overexpressing *tmaR* with a frameshift mutation near the beginning of the gene (**Fig. S3I**, left panel and insert). Hence, increasing TmaR concentration affected not only its pattern of distribution in the cell, but also the cell morphology, causing a severe defect in cell division.

Staining cells overexpressing TmaR-YFP with the DNA binding dye DAPI showed that overexpression of TmaR inhibits cell division after the segregation of the chromosome into each chained cell has occurred (**Fig. 3G**). Staining these cells with FM4-64, an inner-membrane stain (*26*), indicates that constriction of the inner membrane has started, but the extension of TmaR filaments between the chained cells suggest that the process has not been completed (**Fig. 3G**). A closer look at the bridging tubes that connect the cells in the TEM images suggest that in some cases the tubes connecting the cells lack an outer membrane (**Fig. 3F**, lower image).

Together these results confirm that, similar to eukaryotic phase-separated proteins, TmaR can undergo a transition to a less fluid higher structure in the cell, which impairs cell morphology and proliferation.

### TmaR stabilizes flagella-related transcripts at the poles, thus enabling motility and biofilm formation

A significant portion of proteins that form condensates are RNA-binding proteins (*27*). Based on TmaR predicted structure (**Fig. S2B**), the web server BindUP, which employs the NAbind algorithm (*28*), predicts that it has two nucleic acid (NA)-binding regions that come together in space to form an NA-binding domain (**Fig. S4A**, residues in blue in the 3D model and in blue or red in the primary sequence). In support of the importance of the predicted NA-binding domain for TmaR condensation, four out of the six residues that were substituted in our screen for TmaR proteins that are dispersed rather than forming polar condensates (V15, R20, R26, and R91) lie in the predicted NA binding region (**Fig. S4A**, residues in red). We therefore decided to see if RNAs are involved in the process of TmaR condensation by testing the effect of transcriptional arrest on TmaR condensates. The results in **Fig. 4A** show that TmaR condensates dispersed after treating the cells with the transcription inhibitor rifampicin for 10 minutes, a time period longer that the average half-life of *E. coli* transcripts, suggesting that RNAs are important for TmaR phase separation.

**Fig. 4.**
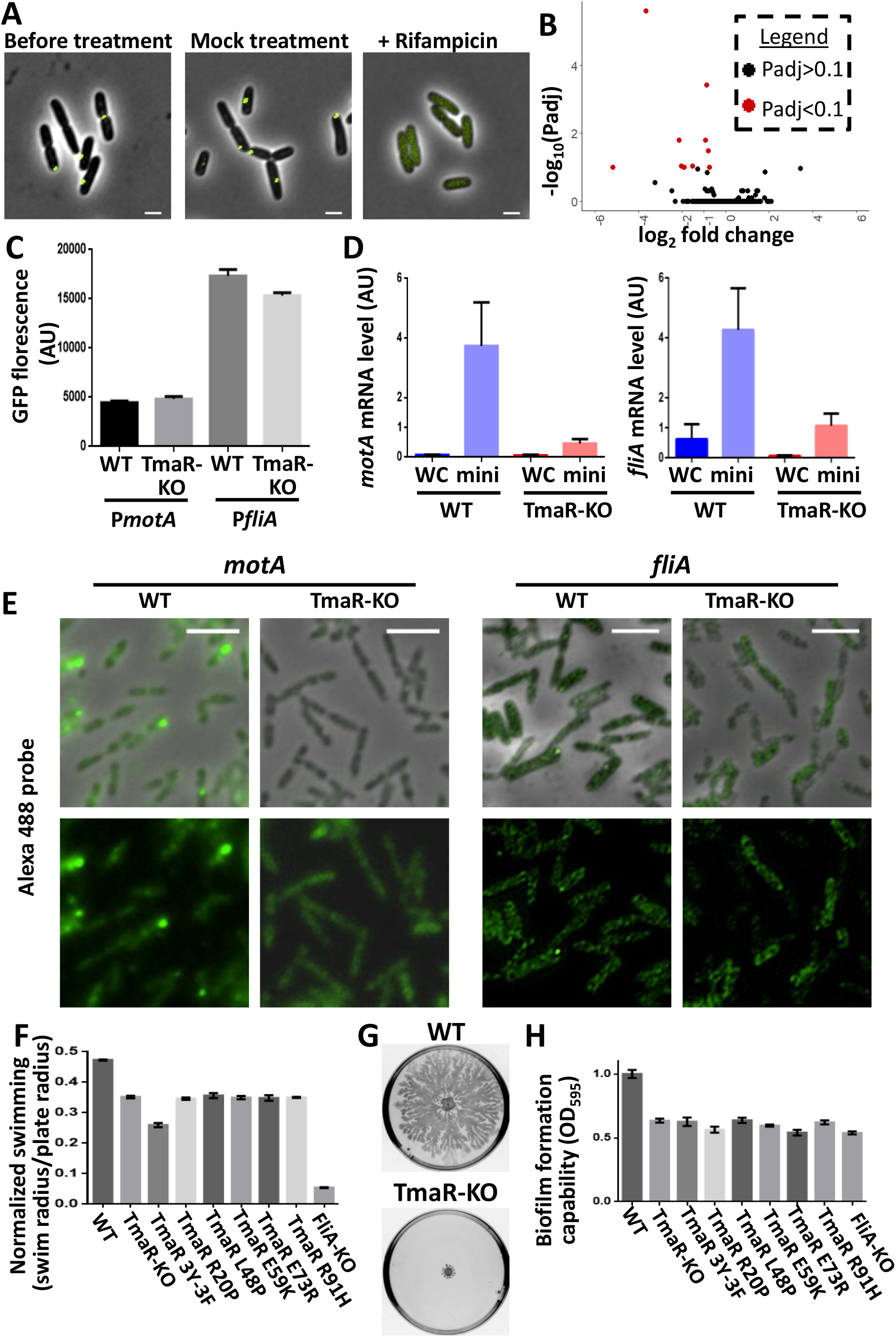
TmaR stabilizes flagella-related transcripts at the poles, thus enabling motility and biofilm formation. (A) Images of cells in mid-logaritmic phase endogenously expressing mYFP-TmaR that were treated for 10 minutes with 2 mg/ml rifampicin dissolved in DMSO or only with DMSO (mock treatment). Scale bar, 2 µm. (B) Volcano plot showing a comparison between the RNA-seq results (log_10_(Padjast)) against Log_2_ fold change) obtained for wild type cells and cells deleted for the *tmaR* gene. Red dots denote the most significantly differentially expressed genes (Padjast<0.1). (C) Average GFP fluorescence in arbitrary units (AU) in wild type cells (WT) and in Δ*tmaR* cells (TmaR-KO) expressing the *gfp* gene from the promoter of *fliA* (P*fliA*) or of *motA* (P*motA*). The bars show the SD between three independent repeats. (D) Bar plots showing *motA* (left) and *fliA* (right) mRNA levels in wild type (WT) or Δ*tmaR* cells (TmaR-KO). RNA was extracted from whole cells (WC) and minicells (mini), the latter representing only the polar content. The *motA* and *fliA* mRNAs were amplified by qPCR and normalized to *gyrA*. (E) *motA* and *fliA* transcripts detected by FISH in wild type (WT) or Δ*tmaR* cells (TmaR-KO). Scale bar, 5 µM. (F) A bar plot showing the normalized swimming (swim radius/plate radius) of wild type cells (WT), Δ*tmaR* cells (TmaR-KO), cells expressing the indicated TmaR mutants or Δ*fliA* cells (FliA-KO). The bars show the SD between three independent repeats. (G) Swarming motility patterns formed by wild type cells (WT) or Δ*tmaR* cells (TmaR-KO) on a semisolid Difco agar plate after incubation for 24 h at 37°C. For swarming of the different TmaR mutants that cannot undergo LLPS, see Fig. S4E. (H) Biofilm formation by wild type cells (WT), Δ*tmaR* cells (TmaR-KO), cells expressing the indicated TmaR mutants or Δ*fliA* cells (FliA-KO) assessed by the crystal violet coloration assay. The bars show the SD between three replicates of three biological repeats.

In light of the reports that LLPS of proteins can facilitate localization of their bound RNAs (*29, 30*), we asked if TmaR might influence RNA localization and/or level. To monitor putative effects on RNA level, we compared the transcriptome of cells expressing wild type TmaR (WT) and cells deleted for the *tmaR* gene (TmaR-KO). Since TmaR is not predicted to be a transcription factor, we did not expect many differences between the strains. Indeed, the level of only 10 transcripts exhibited a significant change between the strains (Padjust<0.1), all showing a decrease by ∼2 fold or higher in TmaR-KO compared to WT cells (**Fig. 4B, Table S1**). Remarkably, 5 out these 10 transcripts are motility-related, mainly flagellar genes: *motA, fliA, flgK, flgA and fliC* (*31*) (**Fig. S4B**). Four of these five motility-related transcripts were previously shown by us to be enriched at the poles of *E. coli* cells (*3*). This, together with the finding reported above, which suggest that RNAs contribute to the formation of the phase-separated TmaR condensates at the poles, intrigued us to ask whether the transcripts whose level is significantly reduced in cells lacking TmaR are part of TmaR condensates.

The level of *motA* and *fliA* transcripts, the first encoding for a major regulator of motility genes (minor sigma factor) and the second for a structural flagella component (*32*–*35*), were affected the most by *tmaR* knockout (35.9 and 12.5 fold reduction, respectively, see **Table S1**). We therefore checked first if the reduction in their level is due to their lower transcription in TmaR-KO cells. The results in **Fig. 4C** show that expression levels of *gfp* from their corresponding promoters [for the library of transcriptional fusions see (*36*)] are quite comparable in cells expressing or lacking TmaR. These results suggest that the reduced levels of *motA* and *fliA* transcripts in TmaR-KO cells in not due to reduced expression, but rather due to their reduced stability.

Next, we aimed at finding out whether TmaR stabilizes these mRNAs. Our assumption was that if the *motA* and *fliA* mRNAs are included in TmaR polar condensates, then not only should they be enriched at the poles, but they are also expected to be absent from the poles of cells lacking TmaR. To check that, we made use of χ1488 *E. coli* strain, which divides asymmetrically near the poles at a higher frequency (*37, 38*), thus generating minicells that package the polar content (*39*). Comparing the level of *motA* and *fliA* extracted from the mini and whole wild type and TmaR-KO cells by RT-qPCR reinforced our previous finding that these transcripts are enriched at the poles (**Fig. 4D**, compare the level in mini and whole cell for *motA*, left panel, and *fliA*, right panel) and demonstrated that their polar enrichment is dramatically reduced at the poles of cells lacking TmaR (**Fig. 4D**, compare the level of each transcript in minicells generated from WT and TmaR-KO cells). Of note, the results were normalized to the equally distributed *gyrA* mRNA, and hence the values document their enrichment, rather than absolute values, explaining the low values of both transcripts level in whole cells. Still, it is clear that the level of *motA* and *fliA* dramatically decreases in TmaR-KO cells compared to wild type (**Fig. 4D**, compare the level of each transcript in WT and TmaR-KO cells), suggesting that their stability sharply drops when not localized to the poles by TmaR.

To directly visualize the subcellular distribution of *motA* and *fliA* transcripts, we used fluorescence in situ hybridization (FISH). Hybridizing fluorescently-labeled probes complementary to either *motA* or *fliA*, we observed *motA* mRNA at the poles, as well as preferred localization of *fliA* to the poles in wild type, but not in TmaR-KO cells (**Fig. 4E**; see quantification in **Fig. S4C**). Together, the results in **Fig. 4D** and **4E** indicate that enrichment of *motA* and *fliA* mRNAs at the cell poles depends on TmaR.

We next asked whether the TmaR-dependent enrichment in motility-related transcripts at the poles is important for the function of their encoded proteins in promoting motility. To answer this question, we compared the ability of wild type cells, TmaR-KO cells and the various TmaR mutants, impaired in forming polar condensates, to perform the two types of flagella-driven motility, i.e., swimming and swarming, the first referring to movement of individual cells in liquid or soft medium and the second to movement on semisolid medium in a synchronized way. The results in **Fig. S4D**, quantified in **Fig. 4F**, show that the swimming ability of TmaR-KO cells and of all the cells expressing TmaR mutants that fail to undergo LLPS is reduced compared to the swimming ability of wild type cells. Moreover, TmaR-KO cells and all TmaR mutants are completely impaired in swarming on semisolid agar **(Fig. 4G** and **S4E**). Of note, the ability or TmaR-KO and all TmaR mutant cells to swim and swarm was comparable to that of cells deleted for the *fliA* gene (FliA-KO), which served as a negative control (*31*). These results imply that not only is TmaR required for swimming and swarming of *E. coli* cells, but its condensation is important for both activities.

The swimming ability of cells expressing a TmaR mutant that cannot get phosphorylated on tyrosine 72 (termed henceforth TmaR 3Y-3F) and does not form polar clusters (*13*) is even lower than that of cells lacking TmaR or expressing the other TmaR mutants, whereas its swarming ability is completely inhibited (**Fig. S4D, 4F, 4G and S4E**). Notably, replacement of E73, a negative amino acid residing next to the phosphorylated tyrosine in a negative patch of TmaR (see **Fig. S2C**), by an amino acid with the opposite charge (E73R) also prevented condensates formation (**Fig. 2C**) and was also defective in swimming and swarming (**Fig. S4D, 4F, 4G and S4E**), albeit to a lesser extent than TmaR 3Y-3F. Hence, we conclude that together the change of charge and the prevention of phosphorylation of this patch account for the observed functional defects of TmaR 3Y-3F.

Since motility is a prerequisite for the initiation of biofilm formation (*40*), we checked the ability of the different strains to form biofilm by the crystal violet colorimetric assay (*41*). The results in **Fig. 4H** show that, as opposed to the ability of wild type cells to form biofilm after 8 hours, cells deleted for TmaR (TmaR-KO) or expressing TmaR mutants that do not form condensates have a dramatically reduced ability to form biofilms, which is slightly higher than that of FliA-KO cells that served as a negative control. Hence, not only motility, but also capabilities that require movement are impaired is cells lacking TmaR or expressing a TmaR mutants that fail to undergo LLPS.

Together our results show that TmaR enables major survival strategies of *E. coli* - motility and biofilm formation - by stabilizing flagella-related transcripts within the polar condensates.

## DISCUSSION

Formation of membraneless micro-scale compartments by LLPS is a newly discovered phenomenon used by cells to organize their content (*6, 42*). LLPS-driven condensates show selective permeability, thus concentrating processes to discreet subcellular regions (*42*). Hence, they are important for bacteria, which are generally devoid of membrane-bound organelles, to gain spatial complexity in their cytoplasm. The existence of condensates in bacteria has been reported (see Introduction), but more evidence for their direct relevance to bacterial physiology and survival is required. Here we show that TmaR, a novel pole localizer, previously shown to control sugar metabolism by spatial regulation of EI activity, forms phase-separated condensates that are important for major bacterial survival capabilities, such as proper division, motility and biofilm formation. Our previous report that TmaR limits the degree of heterogeneity in EI activity by polar sequestration (*13*) is in line with this hypothesis, since phase separation is emerging as a mechanism to effectively reduce variability in protein concentration outside the condensates (*43*). We further show reciprocal relations between TmaR and RNAs. Whereas RNA molecules contribute to phase-separation of TmaR, TmaR recruits flagella-related transcripts to polar condensates, thus stabilizing them. Remarkably, by encompassing these transcripts in the condensates, TmaR enables proper motility and biofilm formation, which are among the most regulated functions (*44*) and regarded as highly important for *E. coli* survival. Of note, accumulating evidence suggest that many stages of the RNA life cycle in eukaryotes occur within biomolecular condensates (*27*). Further research is required for defining the protein and RNA content of TmaR condensates and the interactions among its constituents that enable the control of important pathways, sugar metabolism on one hand, as previously shown by us (*13*), and motility and biofilm formation on the other hand, as shown here.

Emerging evidence suggests different types of coupling between metabolic functions, motility and regulation of flagella assembly, either by direct interaction or via regulatory effects. Examples include activation/regulation of both metabolic functions and bacterial motility by the non-canonical sensory mechanism, by moonlighting proteins, e.g., the citrate cycle enzyme aconitase (AcnB), and by proteins that integrate environmental signals into regulatory circuitry and adaptation, such as the soluble taxis receptor AerC from *A. brasilense*, which dynamically localizes to the cell poles depending on the levels of oxygen (*45*). Specifically, sugar metabolism is linked to motility via the sugar governing system PTS (whose key factor EI is spatially regulated by TmaR), which is involved in regulating motility and taxis (reviewed in (*45*)), and the carbon storage regulator CsrA, which is necessary for motility under various growth conditions and for regulating expression of the master operon for flagellum biosynthesis (*46*). Moreover, CsrA also regulates biofilm formation and dispersal (*47*). Our findings showing that TmaR polar condensates control sugar metabolism, as well as motility and biofilm formation suggest that co-regulation of these major functions is achieved by placing their control centers - the principal PTS protein EI and the flagella-related transcripts - in the same membraneless organelle. This explains the actual “decision making”, as this “organelle” co-regulates sensing and consumption of sugars and produces the appropriate response, that is, swimming away or forming a biofilm. Furthermore, our results add a spatial layer to this mechanism, that is, the cell poles are the arena where co-regulation of all these functions takes place. We previously termed the poles microbrains, since various sensing and signaling pathways localize to them (*48*). The results reported here add RNA stabilization as a rational for assembling transcripts. Hence, although the flagella are randomly distributed around the *E. coli* cell with no apparent concentration at the poles (*49*), placing the flagellar RNAs in the polar membraneless organelle ensures their stabilization and proper expression. This resonates with our demonstration that the chemotaxis-related mRNAs localize to the *E. coli* cell poles, although they encode for both polar and non-polar proteins (*3*). Hence, bacteria seem to invest resources in assembling transcripts for the sake of co-regulation.

It is clear that well established motifs underly much of the activity of TmaR. Starting with unstructured and RNA-binding domains, shown to drive condensation of many proteins (*17*), we show that both the disordered and the ordered domains in TmaR, the latter containing the RNA-binding domain, are required for its LLPS. Recently, condensation of PopZ in *C. crescentus* has been found to be driven by a structured domain, whereas the state of matter of these properties is determined by PopZ disordered region (*50*). Other results presented here show compliance with the molecular grammar developed for LLPS, such as the importance of arginines, often present in RNA-binding domains, for condensation (*20*). Together, our findings suggest that the rules for condensates formation, which may vary from one protein to another, are by and large the same for prokaryotes and eukaryotes.

As opposed to formation of phase-separated condensates, which emerges as an important principle that supports life, formation of aberrant condensates by liquid-to-solid transition is associated with a wide range of human diseases, including neurodegeneration, tumorigenesis, cancer and infectious diseases (*23, 25, 51, 52*). Our results show that bacterial condensates can undergo a similar transition due to the same reasons, i.e., mutations or a change in a protein level. Similar to eukaryotes, a liquid-to-solid transition might have different manifestations and physiological ramifications, depending on what caused it. Hence, an increase in TmaR cellular level leads to the formation of long filaments that can extend from cell to cell and impair bacterial morphology and proliferation, whereas a TmaR variant harboring the E59K point mutation forms aggregates, which are similar in size to the polar condensates formed by wild type TmaR, and impairs cell morphology and division to a lesser extent. The finding that liquid-to-solid transitions that generate aberrant condensates occur also in prokaryotes provides an opportunity to study this diseases-associated process in a simple model system, which might accelerate the development of future therapeutic interventions.

Finally, flagellar motility of *E. coli* cells plays an important role in the development of infections, due to the role of motility in host colonization and biofilm formation, the latter conferring protection against the host immune system, antibiotics, and some environmental stress factors. Hence, attenuation of flagellar motility is regarded as a potential therapy to control *E. coli* pathogenicity (*44*). Discovering a new mechanism for regulating *E. coli* motility is, therefore, of importance.

## ACKNOWLEDGMENTS

We are grateful to Yael Friedmann and Zakhariya Manevitch from the interdepartmental equipment units at the Hebrew University for help with TEM and FRAP microscopy, respectively. We thank Fouad Hassouna and Jamal Fahoum from the Weiner lab for help with adjusting the protocol for protein purification to TmaR. We thank Amirhossein Ghanbari Niaki and Kevin Rhine from Myong Lab for the knowledge acquired during the Physics of the Living Cell Summer School on *in-vitro* studies of LLPS, which took place University of Illinois Urbana-Champaign. We thank Amir Fadel for help with the microscopy screens. We thank Meta Heidenreich, Emmanuel Levy, Ora Furman-Schueler, Martine Ruer, Adam Kłosin and members of the Amster-Choder lab for fruitful discussions and suggestions. Research in OAC lab was supported by the Israel Science Foundation (ISF) founded by the Israel Academy of Sciences and Humanities (grant no. 1274/19). OAC is an incumbent of the Dr. Jacob Grunbaum Chair in Medical Sciences. Research in the MS lab was supported by a VolksWagen (VW) foundation LIFE grant 93092. MS is an incumbent of the Dr. Gilbert Omenn and Martha Darling Professorial Chair in Molecular Genetics

